# Computational Discovery of Evolutionary Conserved Enterovirus 71 RNA Secondary Structures

**DOI:** 10.1101/495507

**Authors:** Nathan Fridlyand, Alexander N. Lukashev, Andrey Chursov, Franco M. Venanzi, Jonathan W. Yewdell, Albert A. Sufianov, Alexander Shneider

## Abstract

Viral RNAs store information in their sequence and structure. Little, however is known about the contribution of RNA structure to viral replication and evolution. We previously described RNA ISRAEU to computationally identify structured regions in viral RNA based on evolutionary conservation of predicted structures. Here, we apply RNA GUESS (Generator of Uniformly Evolved Secondary Structures) to better understand enterovirus 71, a highly pathogenic human picornavirus. RNA GUESS identified previously known evolutionarily conserved picornavirus RNA structures, and uniquely predicted RNA structures with significant structural conservation despite the lack of an obvious sequence conservation. One structure consists of a stretch of obligatorily unpaired nucleotides that can potentially interact with RNA-binding proteins and siRNAs. Another, forms a potential RNA switch that can form two alternative structures while the third and fourth constitute hairpin-like structures. These findings provide a launching point for physical and genetic studies regarding the function and structures of these RNA elements.

## Introduction

RNA secondary structures perform a wide range of important biological functions in addition to their central roles in ribosomes and tRNAs. Auxiliary structural functions include regulating gene expression levels in mammalian cells (2), constraining mutations during bacterial evolution (7), controlling temperature sensitivity (3, 12, 36) and viral replication (1, 34, 35), regulating folding and post-translational modification of encoded proteins (14, 15).

*In silico* prediction of secondary RNA structures is challenging since any given RNA sequence can typically form multiple RNA structures with similar minimal free energies (37). Indeed, highly homologous RNAs can have dissimilar structures (38), while dissimilar sequences can fold into similar structures. An additional difficulty in predicting RNA secondary structure in positive strand RNA viruses is the ubiquitous presence of non-specific structural elements termed Genome-Ordered RNA Structure (GORS) (43).

To improve viral RNA structural predictions, Chursov et al. proposed a new definition of structured RNA regions and a novel computational method RNA ISRAEU to find them (16). Previously, “structured” regions were limited to consecutive nucleotides paired via Watson-Crick (WC) bonds to create stems (40-42). Chursov broadened the definition to include loops and bistable conformations, if they too were evolutionarily conserved across a broad range of strains, reasoning that evolutionary selection implies functional significance. Although the method identifies structurally conserved RNA regions, it did not attempt to characterize the RNA structural elements themselves.

Here, we describe a method to identify specific structures in regions identified by RNA ISRAEU and apply this method to the emerging picornavirus, enterovirus 71 (EV71), an important human pathogen that first appeared in 1969. EV71 is endemic in pediatric populations and can cause devastating epidemics with hundreds of deaths (23-27). We chose EV71 based on the large and diverse public sequence dataset available and the presence of known structural RNA elements that enable validation of our computational methods.

## Results

We applied RNA ISRAEU with minor modifications (described in Materials and Methods) to identify a non-redundant EV71 sequence dataset of representative sequences, as well as EV71 for structured region candidates. The EV71 genome is about 7.5Kb bases, with c. 740 nt. 5’UTR, 6600 nt single open reading frame and about 150 nt 3’UTR. Due to current algorithm limitations, only gapless sequences were used; therefore, the highly structured UTRs of variable length had to be omitted. To establish a length threshold for meaningful analysis of the structured regions identified, we tested the standard deviation moving average for autocorrelation (Supplementary Figure 1.). We chose the second minimum to establish a threshold of 15 or more residues. The structurally-conserved RNA in segments in the EV71 so selected for further analysis are listed in Supplementary Table 1.

In selected regions we identified CSSs (Consistent Structured Sub-regions) as segments of consecutive positions where either all positions inside the segment were paired (WC pairs as well as, canonical RNA, G-U pairs) with exceptionally high or low probabilities. We analyzed RNAfold (17) output to find segments complementary in sequence to the high probability CSSs we identified and thereby generated putative RNA structures. We identified four hairpin-like structures (see below as S1, S2, S3 and S4) containing a stem and a loop between the two sides of the stem. We define the loop positions as positions between the two sides of stem for which the mean probability to be paired is below *P*_*unpaired*_. We also identified a stretch of accessible unpaired nucleotides (S5) and an RNA switch system (S6 and S6-binding regions). All inferred structures were statistically significant (see Statistical Analysis in Methods and Material section for detail).

### Algorithm Validation

The largest structured region, located between positions S1: 4406-4448 contains a physically defined RNA structure (45) (all positions here and below refer to EV71 strain BrCr genome, Genbank U22521). We identified the following high pairing probability CSSs inside S1, S1.1: 4409-4420, and S1.2: 4435-4447. S1.1 sequence matches S1.2 sequence throughout the dataset. While the sequence in positions 4409-4414 matches the sequences in positions 4442-4447, and the sequence in positions 4416-4420 matches the sequence in positions 4435-4439, position 4415 matches position 4440 in some strains and position 4441 in others, which implies that S1 structures in these strains are not identical (Figure 1.a). The S1 structure is the previously identified cis-acting RNA element (cre), a stem-loop structure that templates U addition to the VPg peptide to enable genome replication (28).

**Figure 1.:**
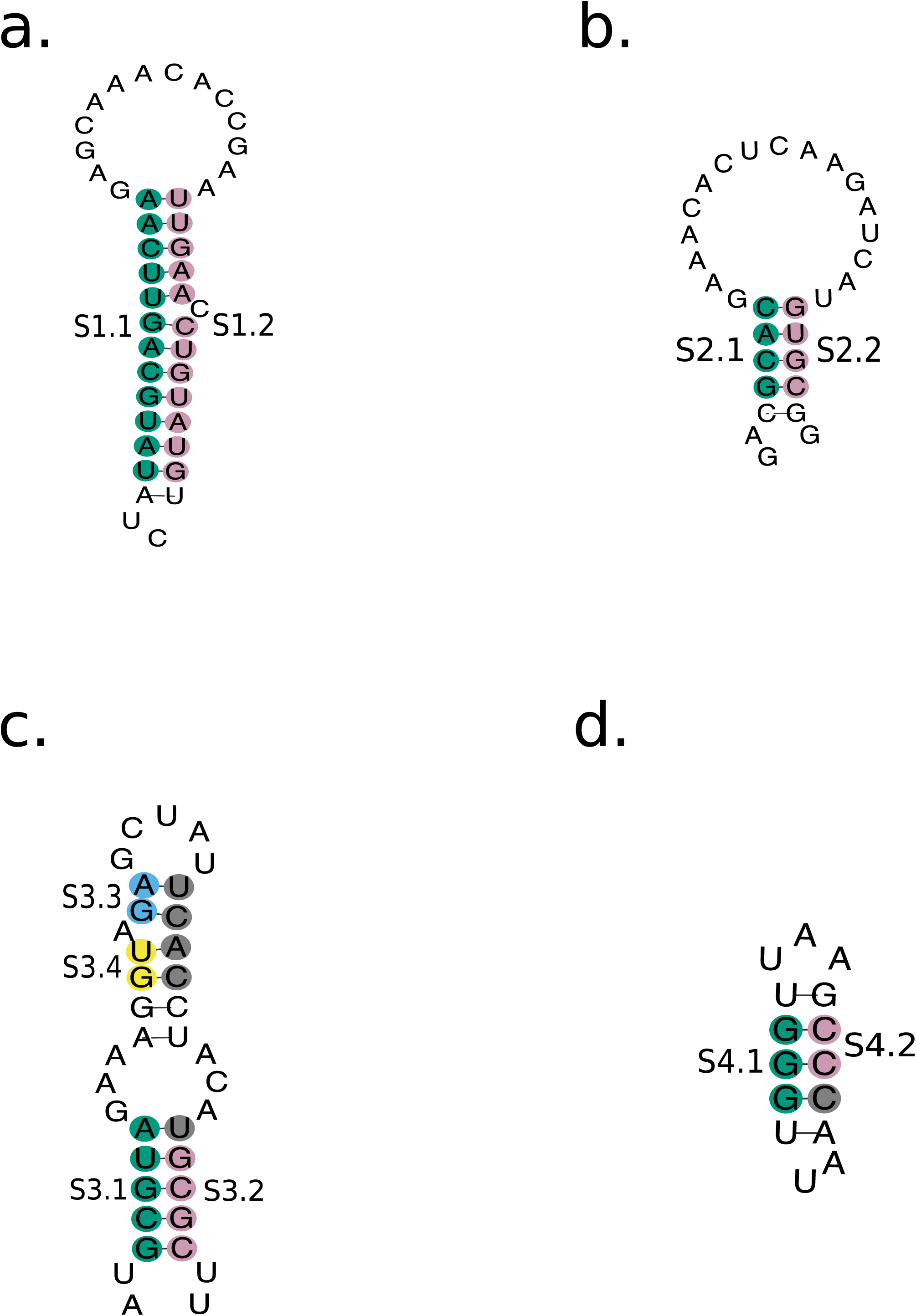
Predicted structures (a) S1, (b) S2, (c) S3 and (d) S4. All present a hairpin formation. Colored are the corresponding CSSs of each structure.

The second largest structured region of the Enterovirus 71 identified by RNA ISRAEU is S2: 7155-7194, which contains two high pairing probability CSSs, S2.1: 7158-7161 and S2.2: 7179-7182. RNA_GUESS reveals that S2.1 and S2.2 have a high tendency to form a double stranded conformation (Figure 1.b) that corresponds to the physically defined α element in poliovirus, a sister species Enterovirus, at positions 6920–7369 (29).

Thus, RNA GUESS successfully identified two physically validated picornavirus RNA structures.

### Newly imputed structures

The third longest structured region identified by RNA ISRAEU was in positions S3’: 2793-2810. RNA GUESS deduced a double stranded conformation with the neighboring structured region S3’’: 2825-2830. Thus, the entire segment S3: 2793-2830 should be treated as single structured region. Two of 4 CSSs in S3 are located at the extremities of S3 and have a high tendency to form a stem with S3.1: 2795-2799 and S3.2: 2825-2828. We also found two CSSs containing 2 nucleotides each in positions S3.3: 2805-2806 and S3.4: 2808-2809. While the very strong tendency of S3.3 and S3.4 to be paired was evident, the identity of their pairing counterparts was more ambiguous. The most common matches (though not unique) for both S3.3 and S3.4 CSSs resided inside the range 2814-2820, with the S3.3 counterpart of highest probability being S3.3p: 2717-2818 and S3.4 counterpart of highest probability being S3.4p: 2815-2816 (Figure 1.c). Base pairing partners for S3.3 and S3.4, segments S3.3P and S3.4P, reside outside of any structured region defined by RNA ISRAEU. Therefore, the statistical prediction was not fully concordant with the intuitively obvious structure, at least not in all inferred folding repetitions.

The next largest structured region was found in positions S4: 5949-5963. The CSSs S4.1: 5950-5952 and S4.2: 5958-5959 possess complementary sequences with an exceptional tendency to form a hairpin structure (Figure 1.d). This structured region was not sequence conserved in the non-redundant dataset, nor did the structure maintains exact pairs in the stem or the exact size of the loop. Yet, in all 94 sequences queried, a predicted stem loop was present.

### Accessible regions

The predicted list of significantly (un)structured RNA regions contains 2 regions constituted of unpaired nucleotides, S5.1: 816-825 and S5.2: 831-840. Due to the proximity of S5.1 and S5.2, as well as the borderline mean and standard deviation values (all positions between 826 and 830 were close to being labeled “unpaired” and close to being labeled structurally-conserved positions), we concluded that the entire region S5: 816-840, should be regarded as a single unpaired region.

Interestingly, this region does not contain G residues in any of the EV71 genomes. Although six of eight codons in the region (ATH for Ile, AAY for Asn, TAY for Tyr) cannot have G in a synonymous position, two codons (ACN for Thr) can have G in the third position. However, none of the EV71 sequences actually have G in the third-position, suggesting strong negative selection. The S5 Sequence Logo (44) is presented in Supplementary figure 2.

### A possible functional switch

The next structured region is S6: 3104-3118. It consists of two CSSs with high probability of WC pairing, S6.1: 3106-3108 and S6.2: 3112-3115. Intriguingly the pairing partners of S6 represent a “three hands handshake” where the middle hand, S6 can pair with S6.L or S6.R (Figure 2). The “right hand”, residing in positions S6.R: 3125-3139 can pair with S6 via S6.3R: 3128-3131 forming bonds with S6.2, and S6.4R: 3137-3139 pairing with S6.1. In some strains S6.4R is located in nt 3136-3138, in others it is located in nt 3135-3137 and in a small number of strains S6.4R resides in 3134-3136. However, this sliding of S6.4R does not modify the formed structures.

**Figure 2.:**
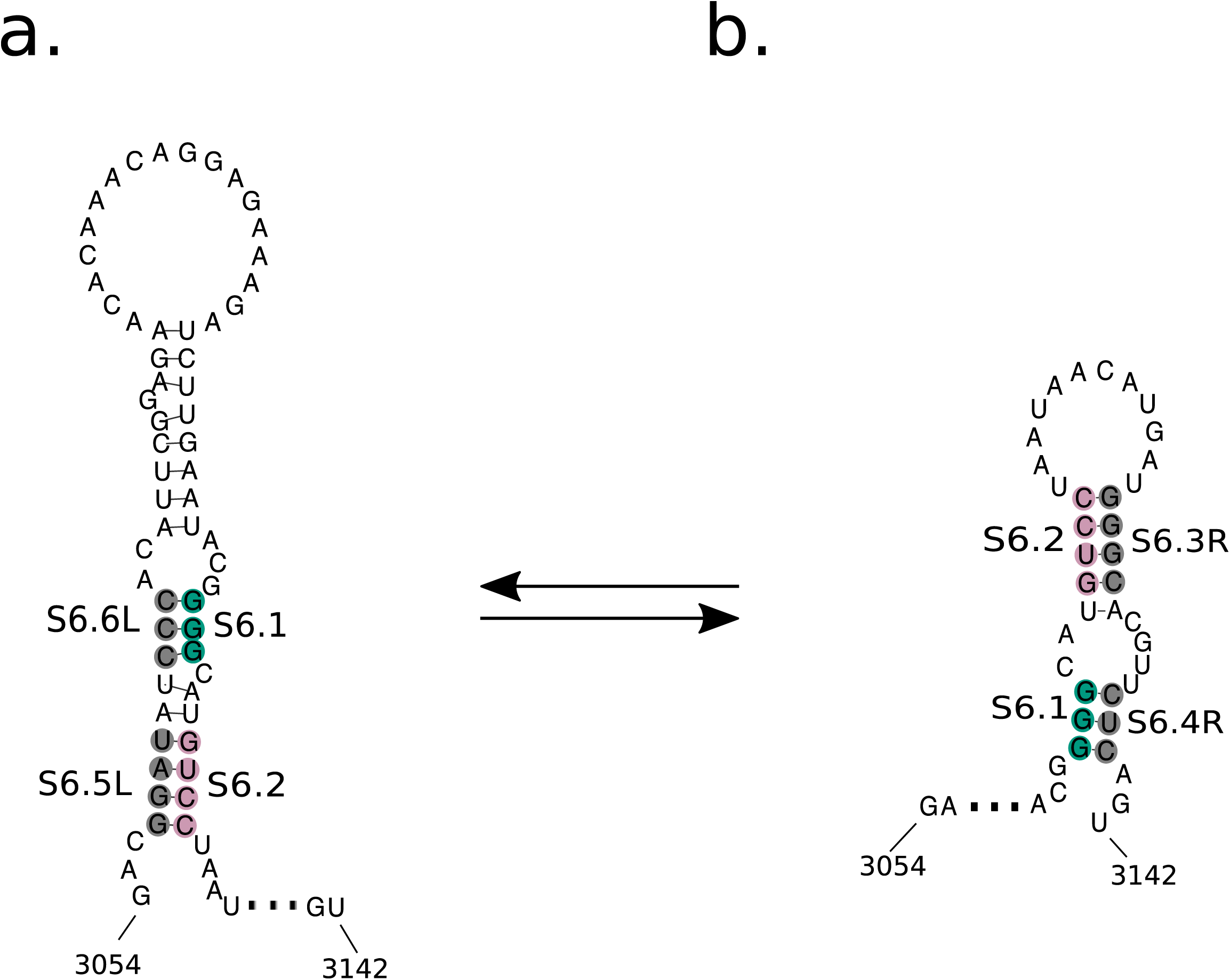
Predicted switch structure S6 with alternating pairings of the high pairing probability CSSs. (a) The CSSs are paired to the left. (b) The CSSs are paired to the right.

The “left hand” resides at positions S6.L: 3054-3067. The pairing of S6 with S6.L is composed of two simultaneous pairings of two elements. Element S6.5L: 3057-3060 is pairing with S6.2, while element S6.6L: 3063-3065 is pairing with S6.1 (Figure 2).

Notably, S6.3R, S6.4R, S6.5L and S6.6L are not CSSs because they are located outside of structured RNA regions. Rather, they are partners of structurally conserved CSSs. Since only two of the three hands can shake at any one time, this structure can function as an RNA switch.

## Discussion

Recently, we proposed a new definition of structured RNA regions and devised RNA ISRAEU to identify these regions (16). Here, we extend our original findings on influenza A virus to EV71, which contains a highly conserved physically defined structure, which was correctly identified by RNA GUESS. RNA GUESS extends the functionality of RNA ISRAEU by predicting evolutionarily conserved structures from RNAfold pairing probability predictions. RNA GUESS does not identify all secondary RNA structures, as it is limited to structures preserved throughout the sequence dataset. While unable to provide an exhaustive set of the secondary RNA structures, RNA GUESS is unlikely to generate false-positives. Thus, although our method is limited to evolutionarily conserved structures, this limitation increases it accuracy. The power of RNA GUESS is highlighted by identification of a structure in EV71 that was recently reported for poliovirus, a distantly related enterovirus. This structure contains just four base pairs, and could be easily overlooked among numerous structures that RNAfold can predict in a single sequence.

Secondary RNA structures may be entirely formed by the nucleotides of the structured region, as with the S4 hairpin structure which is present in every EV71 strain in the dataset. Alternatively, a structured RNA region may contain half of a stem whose other half may be constituted by nucleotides outside of the structured region, with positions that vary with viral strain. This is analogous to a belt buckle that can attach to different holes, and neatly describes the S6 structure of EV71 mRNA. Again, such prediction is highly unlikely be made by analyzing a single sequence.

RNA can form alternative structures with similar minimal free energies. We provide evidence that EV71 may contain a RNA switch composed of an RNA element, S6, that is always paired with two potential up or downstream pairing elements. If confirmed experimentally, this would be the first RNA switch discovered in enteroviruses. It is of obvious interest to determine the functional significance of this putative switch, factors that controls the switching (e.g. other RNAs, ions, etc.), and how common such switches are in other viruses.

We show here that RNA structural conservation may also consist of unpaired bases, as in the element S5. As S5 resides at the beginning of the sole enterovirus ORF, we hypothesize that this region may constitute a binding site for external regulatory RNA segments or for RNA-binding proteins. We note that the evolutionary importance of an unpaired S5 region is supported by conserved codon bias in the region, since S5 lacks guanines in all strains examined, consistent with maintaining an unpaired structure.

A region lacking G residues would have lower chance of being paired. G forms two pairs with similar energies, G-U and G-C. Moreover, mismatches that contain G have the lowest impact on region’s overall pairing energy (30). S5 region does not contain Guanines, which is not true for EV71 genome as a whole. This could be either due to a pressure to avoid internal bonding within S5 or due to a general phenomenon of CpG dinucleotide avoidance in RNA viruses (31, 32, 33). However, the overall relative CpG content was only 0.52 of the expected for the EV71 alignment we used. Therefore, complete absence of G residues at synonymous positions 827 and 830 in all 94 sequences in the dataset cannot be explained by CpG dinucleotide avoidance only and should come from a pressure to avoid base pair stems within S5. Still, when two factors contribute to the phenomena, it is difficult to quantify statistical significance of each one. Interestingly, Guanine avoidance in this genome region was evident on the level of the whole EV-A species (which includes also EV71). Among all 260 non-EV71 EV-A sequences available in GenBank, this genome region contained just 0.2% G residues, which further supports that it is an obligatorily unpaired (thus accessible) region.

Experimental verification of our *in silico* structural predictions is, of course, essential, using single molecule (18, 19) and/or high throughput techniques (20-22). The functional importance of each structure can be tested by creating recombinant viruses with synonymous substitutions that alter the inferred RNA structure. Phenotypic characterization of the mutants should reveal the contribution of a given RNA structure to the viral replication cycle. By creating mutants with a high number of synonymous mutations it may be possible to develop highly attenuated vaccines strains that maintain high immunogenicity by synthesizing large amounts of viral proteins in infected cells while having severe replication defects.

## Methods and Materials

### Enterovirus71 mRNA data set

Our original dataset included all 524 complete ORFs encoding polyproteins of Enterovirus71 strains which were available in Genbank as of December 2016. (Supplementary Table 2). All EV71 mRNA sequences contained 6579 nucleotides positions were aligned. Entries that contained obvious sequencing errors (e.g. frameshifts or stop codons) and sequences of modified strains were omitted. Wobble positions were resolved to the consensus sequence. Sequences which were at Hamming distance of less than 3% from any other sequence in the alignment were omitted as described in (16) resulting in a non-redundant dataset of 94 sequences (Supplementary Table 3).

### RNAfold

We used the RNAfold software from the Vienna RNA package for the identification of structured RNA regions as well as ensuing structure analysis (4).

The command line we use was:

$ RNAfold -p -T 37 --noLP

The “–p” option produces the partition function and the matrix of pairing probabilities

The “-T 37” option, sets the temperature to 37°C (also the default setting.)

The “--noLP” option restricts the length of created helixes (stems) to be no shorter than 2.

### Identification of structured regions

Structured RNA regions were identified according to Chursov et al. (16). If two structured regions were separated by one or two non structurally-conserved positions, we fused them into a single structured region. Also, structured region S3’, in positions 2793-2810, had a very strong tendency to be paired with closely located short structured region in positions 2825-2830. Thus, we fused these two regions into one, S3. Finally, we fused together structured regions S5.1 and S5.2 forming S5. Both S5.1 and S5.2 were identified as unpaired regions separated just by 5 nucleotides. All 5 nucleotides were on the verge of being qualified as “structure-conserved” according to (16).

The method for defining structured regions as it was presented in (16) has generated regions of different length. Obviously, EV71 mRNA is much longer than influenza mRNAs which were used to develop and validate the method (16). Thus, for EV71, the method can generate some short nucleotide stretches which could be false positive. Analysis of data autocorrelation allows identifying the minimal length below which the risk of false positive is substantial. We built a profile of standard deviation moving averages as described in (16) and identified the autocorrelation in the data (Supplementary Figure 1). The threshold is bounded by the first minimum (Lag = 6) and the second minimum (Lag = 16). It was established to be 15, i.e., any structured region shorter than 15 was at high risk of being a false positive. Thus, we only analyzed conserved regions no shorter than 15 nucleotides.

### Defining paired and unpaired nucleotides

We analyzed experimentally confirmed RNA structures and found that according to RNAfold their stems are constituted of nucleotides with probability to be paired above *P*_*paired*_ == 0.85. The paring probabilities of the nucleotides in the loops were below *P*_*unpaired*_ == 0.274. Thus, we call nucleotide positions with pairing probabilities of 0.85 or higher as “paired” and 0.274 and lower as “unpaired”. Noteworthy, all nucleotides within S5.1 and S5.2 are unpaired; and all 5 nucleotides in the linker between S5.1 and S5.2 are close to being unpaired with their pairing probabilities being between 0.287 and 0.328. Thus, we consider the entire S5 as an unpaired region.

### Identifying evolutionary conserved structures and switches within structured regions

#### Evolutionary conserved structures

We define high pairing probability CSSs as the longest possible segments of consecutive “paired” positions inside a structured region. We build RNA structure(s) for an RNA structured region by identifying CSSs within the region, identifying RNA segments which are evolutionary conserved coupling partners for the CSSs, and linking them together. We define evolutionary conserved CSS coupling partners via a sequence of three steps.

Step 1: Identify regions within the RNA molecule, which are complementary to the CSS, as

1. The length of the complementary region is equal to the length of the CSS.
2. Each nucleotide within the CSS is WC paired to the corresponding nucleotide within the complementary region.
3. Probability of each WC paired couple within the region (where one nucleotide belongs to the CSS and the nucleotide it is coupled with belongs to the complementary region) is +/-10% of the mean paring probability for the region. For example, if the mean probability for a nucleotide of the CSS to be coupled with a counterpart from the CSS-complementary region is 0.6, no pair within the region would have its pairing probability outside of 0.54-0.66 range (supplementary Figure 3).

Step 2: Select prevalent CSS-pairing regions. A pair of CSS and its CSS-complementary region is considered evolutionary conserved if it can be found in at least 80% of the viral strains of the non-redundant dataset satisfying all three above listed criteria in each strain.

#### Evolutionary conserved functional switch

We defined potential switches similar to identifying CSSs and their evolutionary conserved pairing sequences as described above with two exceptions:

1. CSSs and their binding counterparts had to be present in 50% of the strains in the non-redundant dataset.
2. A CSS should possess at least 2 alternative potential coupling counterparts.

#### Statistical Analysis

We characterized each structure S1-S4 by the longest stem and the longest loop. The longest stem was defined as the longest high pairing probability CSS of the structure. The longest loop was defined as the longest stretch of positions between the two sides of a stem for which the mean probability to be paired was less than *P*_*unpaired*_. Parameters Stem Size and Loop Size for S1-S4 are presented in Table 1.

**Table 1.:** Parameters of statistical analysis for predicted structures S1-S5 are: (1) Region length, (2) Longest consecutive stem of the structure, (3) Maximal amount of loop positions as defined in Methods, (4) Randomly selected, of same length as the predicted structure, segments of the EV71 coding region with stem of Stem Size or larger and loop of Loop Size or larger.

|  | Length | Stem Size | Loop Size | Qualifying Segments | p-value |
| --- | --- | --- | --- | --- | --- |
| **S1** | 43 | 6 | 10 | 3/1000 | <0.0001 |
| **S2** | 40 | 4 | 15 | 0/1000 | <0.0001 |
| **S3** | 38 | 5 | 4 | 5/1000 | <0.0001 |
| **S4** | 15 | 2 | 2 | 27/1000 | <0.0005 |
| **S5** | 25 | NA | 18 | 0/1000 | <0.0001 |

For each structure S1-S4, we have randomly selected 1000 regions of the same length within the EV71 mRNA. We examined the probability of observing a region satisfying all three conditions

1. A random region contained the same or greater number of loop positions as the identified structure.
2. The region contained the same or greater length stem as the identified structure.
3. The stem satisfied the three steps presented at Evolutionary conserved structures section.

Apparently, probability to find regions satisfying all three criteria exceeds and overestimates probability to find the actual structures S1-S4 because it assessed only 2 elements, the longest stem and the number of loop positions, while the actual structure has other elements too (e.g. shorter stems).

For S5, we have randomly selected 1000 regions. We examined the probability of observing a region with the same or greater number of loop positions as in S5.

The null hypothesis (H_0_) was defined as the probability of a region to be ≥ 5%.

P-values were evaluated using one-sided binomial test, with N=1000, K=Number of random regions containing a similar structure and P=0.05.

Rejection of H_0_ suggests that the observed structures are not likely to reside in their corresponding structured regions by chance (12).

We do not believe that it would be correct to apply statistical analysis based only on the lengths of pairing elements to a switch-like structure of S6 because for the situation of two alternative shapes formed by the switch the “diameters” of the loops and other spatial parameters, which we cannot assess based on one structure only, are expected to play a crucial role. For example, is it crucial that the distance between the central and the right pairing partners is 7 nucleotides, and the distance between the central and the left pairing counterpart is 37. Depending on the restrictions on the lengths of the two loops the statistical analysis may differ in the p-value many orders of magnitude. These restrictions are not applicable to the accessible unpaired region of S6 and much less applicable to simple hairpin structures like S1-S4.

## Acknowledgements

We would like to thank Dr. Leonid Kuznetsov for fruitful discussions.

## SUPPLEMENTARY MATERIAL

**Supplementary Figure 1.:** SD moving average autocorrelation. Based on the alignment of the non-redundant set of strains, we calculate the SD moving average by applying a sliding window approach to smooth individual fluctuations of standard deviations of nucleotide base pairing probabilities. The SD moving average autocorrelation demonstrates that only the first two minima (at Lag=6 and Lag=16) should determine the length threshold for structured regions. Structured regions shorter than the threshold are considered indistinguishable from random noise and thus are not further analyzed.

**Supplementary Figure 2.:** S5 Sequence Logo, showing complete avoidance of G, even in positions 827 and 830, where a synonymous mutation with G is possible.

**Supplementary Figure 3.:** (a) A valid evolutionary conserved pairing of regions. The mean paring probability of the 5bp stem is 0.6. All five of the stem’s pairs are of probabilities greater than 0.54 and are less than 0.66. (b) An invalid evolutionary conserved pairing of regions. While the mean pairing probability is 0.8 (for the 5bp stem), the binding probabilities for the third pair (0.9) and for the forth pair (0.65) are outside the 10% margin from the mean.

**Supplementary Table1.:** Structurally conserved RNA regions residing in EV71 coding region. Bold highlighted regions were further analyzed.

| Start Position | End Position | Length |
| --- | --- | --- |
| **4406** | **4448** | **43** |
| **7155** | **7184** | **30** |
| **2793** | **2810** | **18** |
| **3104** | **3118** | **15** |
| **5949** | **5963** | **15** |
| 1093 | 1104 | 12 |
| 3211 | 3222 | 12 |
| **815** | **825** | **11** |
| **830** | **840** | **11** |
| 1158 | 1168 | 11 |
| 1318 | 1328 | 11 |
| 1354 | 1364 | 11 |
| 2847 | 2856 | 10 |
| 2444 | 2452 | 9 |
| 2716 | 2724 | 9 |
| 1122 | 1129 | 8 |
| **2821** | **2828** | **8** |
| 1053 | 1059 | 7 |
| 1177 | 1183 | 7 |
| 2593 | 2599 | 7 |
| 3671 | 3677 | 7 |
| 3721 | 3727 | 7 |
| 5572 | 5578 | 7 |
| 1764 | 1769 | 6 |
| 2240 | 2245 | 6 |
| 2832 | 2837 | 6 |
| 3349 | 3354 | 6 |
| 3660 | 3665 | 6 |
| 4808 | 4813 | 6 |
| 5872 | 5877 | 6 |
| 7144 | 7149 | 6 |
| 7233 | 7238 | 6 |
| 805 | 809 | 5 |
| 853 | 857 | 5 |
| 895 | 899 | 5 |
| 1073 | 1077 | 5 |
| 1267 | 1271 | 5 |
| 1492 | 1496 | 5 |
| 1516 | 1520 | 5 |
| 1585 | 1589 | 5 |
| 1735 | 1739 | 5 |
| 1855 | 1859 | 5 |
| 2103 | 2107 | 5 |
| 2202 | 2206 | 5 |
| 2570 | 2574 | 5 |
| 2784 | 2788 | 5 |
| 2971 | 2975 | 5 |
| 3127 | 3131 | 5 |
| 3227 | 3231 | 5 |
| 3376 | 3380 | 5 |
| 3706 | 3710 | 5 |
| 3752 | 3756 | 5 |
| 4164 | 4168 | 5 |
| 4306 | 4310 | 5 |
| 4683 | 4687 | 5 |
| 4962 | 4966 | 5 |
| 5034 | 5038 | 5 |
| 5282 | 5286 | 5 |
| 5527 | 5531 | 5 |
| 5740 | 5744 | 5 |
| 5903 | 5907 | 5 |
| 6787 | 6791 | 5 |
| 6833 | 6837 | 5 |
| 7022 | 7026 | 5 |
| 7065 | 7069 | 5 |

## References

1. Tuplin A. Diverse roles and interactions of RNA structures during the replication of positive-stranded RNA viruses of humans and animals. J Gen Virol 2015; 96: 1497–1503.

2. Ilyinskii PO, Schmidt T, Lukashev D, Meriin AB, Thoidis G, Frishman D and Shneider AM. Importance of mRNA secondary structural elements for the expression of influenza virus genes. OMICS 2009; 13: 421–430.

3. Chursov A, Kopetzky SJ, Bocharov G, Frishman D and Shneider A. RNAtips: Analysis of temperature-induced changes of RNA secondary structure. Nucleic Acids Res 2013; 41: W486–491.

4. Shabalina SA, Spiridonov NA and Kashina A. Sounds of silence: synonymous nucleotides as a key to biological regulation and complexity. Nucleic Acids Res 2013; 41: 2073–2094.

5. Geisler S and Coller J. RNA in unexpected places: long non-coding RNA functions in diverse cellular contexts. Nat Rev Mol Cell Biol 2013; 14: 699–712.

6. Wan Y, Qu K, Ouyang Z, Kertesz M, Li J, Tibshirani R, Makino DL, Nutter RC, Segal E and Chang HY. Genome-wide measurement of RNA folding energies. Mol Cell 2012; 48: 169–181.

7. Chursov A, Frishman D and Shneider A. Conservation of mRNA secondary structures may filter out mutations in Escherichia coli evolution. Nucleic Acids Res 2013; 41: 7854–7860.

8. Bevilacqua PC and Blose JM. Structures, kinetics, thermodynamics, and biological functions of RNA hairpins. Annu Rev Phys Chem 2008; 59: 79–103.

9. Wan Y, Kertesz M, Spitale RC, Segal E and Chang HY. Understanding the transcriptome through RNA structure. Nat Rev Genet 2011; 12: 641–655.

10. Rouskin S, Zubradt M, Washietl S, Kellis M and Weissman JS. Genome-wide probing of RNA structure reveals active unfolding of mRNA structures in vivo. Nature 2014; 505: 701–705.

11. Mortimer SA, Kidwell MA and Doudna JA. Insights into RNA structure and function from genome-wide studies. Nat Rev Genet 2014; 15: 469–479.

12. Chursov A, Kopetzky SJ, Leshchiner I, Kondofersky I, Theis FJ, Frishman D and Shneider A. Specific temperature-induced perturbations of secondary mRNA structures are associated with the cold-adapted temperature-sensitive phenotype of influenza A virus. RNA Biol 2012; 9: 1266–1274.

13. Dethoff EA, Chugh J, Mustoe AM and Al-Hashimi HM. Functional complexity and regulation through RNA dynamics. Nature 2012; 482: 322–330.

14. Faure G, Ogurtsov AY, Shabalina SA, Koonin EV. Role of mRNA structure in the control of protein folding. Nucleic Acids Res 2016; 44: 10898–10911; doi:10.1093/nar/gkw671.

15. Faure G, Ogurtsov AY, Shabalina SA, Koonin EV. Adaptation of mRNA structure to control protein folding. RNA Biol 2017; 14:1649–1654; doi:10.1080/15476286.2017.1349047

16. Chursov A, Fridlyand N, Sufianov AA, Kiselev OI, Baranovskaya I, Vasin, A, Yewdell JW and Shneider A. Novel Computational Method to Define RNA PSRs Explains Influenza A Virus Nucleotide Conservation. doi:10.1101/494336

17. Lorenz R, Bernhart SH, Honer Zu Siederdissen C, Tafer H, Flamm C, Stadler PF and Hofacker IL. ViennaRNA Package 2.0. Algorithms; Mol Biol 2011; 6. 26

18. Wilkinson KA, Merino EJ, Weeks KM. Selective 2’-hydroxyl acylation analyzed by primer extension (SHAPE): quantitative RNA structure analysis at single nucleotide resolution. Nat Protoc. 2006; 1(3): 1610–6; doi:10.1038/nprot.2006.249

19. Kim SH, Quigley G, Suddath FL, Rich A. High-Resolution X-Ray Diffraction Patterns of Crystalline Transfer RNA That Show Helical Regions. Proc Natl Acad Sci U S A 1971; 86(4): 841–5

20. Kertesz M, Wan Y, Mazor E, Rinn JL, Nutter RC, Chang HY, Segal E. Genome-wide measurement of RNA secondary structure in yeast. Nature 2010; 467(7311):103–7; doi: 10.1038/nature09322

21. Lucks JB, Mortimer SA, Trapnell C, Luo S, Aviran S, Schroth GP, Pachter L, Doudna JA, Arkin AP. Multiplexed RNA structure characterization with selective 2’-hydroxyl acylation analyzed by primer extension sequencing (SHAPE-Seq). Proc Natl Acad Sci U S A 2011; 108(27):11063–8; doi: 10.1073/pnas.1106501108

22. Underwood JG, Uzilov AV, Katzman S, Onodera CS, Mainzer JE, Mathews DH, Lowe TM, Salama SR, Haussler D. FragSeq: transcriptome-wide RNA structure probing using high-throughput sequencing. Nat Methods 2010; 7(12):995–1001; doi: 10.1038/nmeth.1529.

23. McMinn PC. 2012. Recent advances in the molecular epidemiology and control of human enterovirus 71 infection. Curr Opin Virol 2:199–205

24. Bible JM, Iturriza-Gomara M, Megson B, Brown D, Pantelidis P, Earl P, Bendig J, Tong CY. 2008. Molecular epidemiology of human enterovirus 71 in the United Kingdom from 1998 to 2006. J Clin Microbiol 46:3192–200

25. Hassel C, Mirand A, Lukashev A, TerletskaiaLadwig E, Farkas A, Schuffenecker I, Diedrich S, Huemer HP, Archimbaud C, Peigue-Lafeuille H, Henquell C, Bailly JL. 2015. Transmission patterns of human enterovirus 71 to, from and among European countries, 2003 to 2013. Euro Surveill 20:30005

26. Vallet S, Legrand Quillien MC, Dailland T, Podeur G, Gouriou S, Schuffenecker I, Payan C, Marcorelles P. 2009. Fatal case of enterovirus 71 infection, France, 2007. Emerg Infect Dis 15:1837–40

27. Akhmadishina LV, Govorukhina MV, Kovalev EV, Nenadskaya SA, Ivanova OE, Lukashev AN. 2015. Enterovirus A71 Meningoencephalitis Outbreak, Rostov-on-Don, Russia, 2013. Emerg Infect Dis 21:1440–3

28. Steil BP, Barton DJ. 2009. Cis-active RNA elements (CREs) and picornavirus RNA replication. Virus Res 139:240–52

29. Song Y, Liu Y, Ward CB, Mueller S, Futcher B, Skiena S, Paul AV, Wimmer E. 2012. Identification of two functionally redundant RNA elements in the coding sequence of poliovirus using computer-generated design. Proc Natl Acad Sci U S A 109:14301–7

30. Hooyberghs J, Van Hummelen P, Carlon E. The effects of mismatches on hybridization in DNA microarrays: determination of nearest neighbor parameters. Nucleic Acids Res 2009, 37: e53

31. Jenkins GM, Holmes EC. The extent of codon usage bias in human RNA viruses and its evolutionary origin. Virus Res 2003, 92:1-7

32. Belalov IS, Lukashev AN. Causes and implications of codon usage bias in RNA viruses. PLoS One 2013, 8: e56642

33. Takata MA, Goncalves-Caneiro D, Zang TM, Soll SJ, York A, Blanco-Melo D, Bieniasz PD. GC dinucleotide suppression enables antiviral defence targeting non-self RNA. Nature 2017, 550:124–127

34. Spronken MI, van de Sandt CE, de Jongh EP, Vuong O, van der Vliet S, Bestebroer TM, Olsthoorn RCL, Rimmelzwaan GF, Fouchier RAM, Gultyaev AP. A compensatory mutagenesis study of a conserved hairpin in the M gene segment of influenza A virus shows its role in virus replication. RNA Biol. 2017. 14:1606–1616

35. Gultyaev AP, Fouchier RA, Olsthoorn RC. Influenza virus RNA structure: unique and common features. Int Rev Immunol. 2010. 29:533–56

36. Shamovsky I, Ivannikov M, Kandel ES, Gershon D, Nudler E. RNA-mediated response to heat shock in mammalian cells. Nature. 2006. 440:556–60.

37. Micura R, Höbartner C. On secondary structure rearrangements and equilibria of small RNAs. Chembiochem. 2003. 4:1263

38. Chursov A1, Walter MC, Schmidt T, Mironov A, Shneider A, Frishman D. Sequence-structure relationships in yeast mRNAs. Nucleic Acids Res. 2012. 40:956–62

39. Nestor G. Iglesias, Andrea V. Gamarnik. Dynamic RNA structures in the dengue virus genome. RNA Biol. 2011. 8:249–257

40. Seetin MG, Mathews DH. RNA structure prediction: an overview of methods. Methods Mol Biol. 2012. 905:99–122.

41. Gruber AR, Findeiss S, Washietl S, Hofacker IL, Stadler PF. RNAz 2.0: improved noncoding RNA detection. Pac Symp Biocomput. 2010. 69–79.

42. Soldatov RA, Vinogradova SV, Mironov AA. RNASurface: fast and accurate detection of locally optimal potentially structured RNA segments. Bioinformatics. 2014. 30:457–63.

43. Davis M, Sagan SM, Pezacki JP, Evans DJ, and Simmonds P. Bioinformatic and Physical Characterizations of Genome-Scale Ordered RNA Structure in Mammalian RNA Viruses. J. Virol. 2008. 82:11824–11836

44. Crools GE, Hon G, Chandonia JM and Brenner SE. WebLogo: A Sequence Logo Generator. Genome. Res. 2004. 14:1188–1190

45. Goodfellow I, Chaudhry Y, Richardson A, Meredith J, Almond JW, Barclay W and Evans DJ. Identification of a cis-acting replication element within the poliovirus coding region. J. Virol. 2000. 74:4590–600

